# Picture naming yields highly consistent cortical activation patterns: test-retest reliability of magnetoencephalography recordings

**DOI:** 10.1101/2020.10.08.330886

**Authors:** Heidi Ala-Salomäki, Jan Kujala, Mia Liljeström, Riitta Salmelin

**Affiliations:** Department of Neuroscience and Biomedical Engineering, Aalto University, FI-00076 Aalto, Finland; Aalto NeuroImaging, Aalto University, FI-00076 Aalto, Finland; Department of Psychology, University of Jyväskylä, FI-40014 University of Jyväskylä, Finland

**Keywords:** picture naming, MEG, reproducibility, test-retest, semantic judgment, individual assessment

## Abstract

Reliable paradigms and imaging measures of individual-level brain activity are paramount when reaching from group-level research studies to clinical assessment of individual patients. Magnetoencephalography (MEG) provides a direct, non-invasive measure of cortical processing with high spatiotemporal accuracy, and is thus well suited for assessment of functional brain damage in patients with language difficulties. This MEG study aimed to identify, in a picture naming paradigm, source-localized evoked activity and modulations of cortical oscillations that show high test-retest reliability across measurement days in healthy individuals, demonstrating their applicability in clinical settings. For patients with a severe language disorder picture naming can be a challenging task. Therefore, we also determined whether a semantic judgement task (‘is this item living’) with a spoken response (‘yes’/’no’) would suffice to induce comparably consistent activity within brain regions related to language production. The MEG data was collected from 19 healthy participants on two separate days. In picture naming, evoked activity was consistent across measurement days (intraclass correlation coefficient (ICC)>0.4) in the left frontal (400–800 ms after image onset), sensorimotor (200–800 ms), parietal (200–600 ms), temporal (200–800 ms), occipital (400–800 ms) and cingulate (600–800 ms) regions, as well as the right temporal (600–800 ms) region. In the semantic judgement task, consistent evoked activity was spatially more limited, occurring in the left temporal (200–800 ms), sensorimotor (400–800 ms), occipital (400–600 ms) and subparietal (600–800 ms) regions, and the right supramarginal cortex (600–800 ms). The naming task showed typical beta oscillatory suppression in premotor and sensorimotor regions (800–1200 ms) but other consistent modulations of oscillatory activity were mostly observed in posterior cortical regions that have not typically been associated with language processing. The high test-retest consistency of MEG evoked activity in the picture naming task testifies to its applicability in clinical evaluations of language function, as well as in longitudinal MEG studies of language production in clinical and healthy populations.

## 1 Introduction

Language production is a dynamic process that requires integration of information between several brain regions (Liljeström et al., 2015a,b; Simonyan and Fuertinger, 2015), making it highly susceptible to impairment in neurological disorders (Boschi et al., 2017; Price et al., 2010). MEG provides a highly time-resolved, direct measure of neural activity underlying language processing, and thus holds promise for evaluation of language mapping in clinical populations. In post-stroke aphasia, MEG recordings of language tasks have shown potential in tracking treatment-induced recovery processes (Cornelissen et al., 2003) and in identifying perilesional cortical areas as targets for brain-stimulation based individualized rehabilitation protocols (Chu et al., 2015). MEG recordings can also aid in diagnosis and prognosis of patients with primary progressive aphasia (Kielar et al., 2018) and dementias (Pievani et al., 2011). For these purposes, reliable measures of brain activity in individual participants are paramount. MEG evoked and oscillatory activity provide measures at the single-subject level (Golubic et al., 2017; Laaksonen et al., 2012; Liljeström et al., 2009; Salmelin et al., 1994). It is, however, unclear how consistent the individual MEG activation patterns are in various language tasks. The aim of the current study was to identify activation patterns related to preparation for language production that show high test-retest reliability across measurement days in healthy individuals.

Earlier studies on the reliability of language mapping tasks have primarily used fMRI (Eaton et al., 2008; Gorgolewski et al., 2013; Harrington et al., 2006; Meltzer et al., 2009; Morrison et al., 2016; Nettekoven et al., 2018; Rau et al., 2007; Rutten et al., 2002; Wilson et al., 2017). These studies show variable results, with reliability ranging from moderate to high. To date, there is no consensus on consistent neural activations related to language production. For example, activation in the left inferior frontal gyrus (IFG), an area highly frequently linked to successful language production, is not systematically found in every study (Meltzer et al., 2009; Nettekoven et al., 2018; Rau et al., 2007). It has been suggested that poor reliability might be a consequence of, e.g., task habituation and learning (Meltzer et al., 2009; Nettekoven et al., 2018; Rau et al., 2007). The baseline task also seems to have an impact on the results. This is especially relevant in picture naming, as the identified amount of visual object processing depends on the baseline task (Price et al., 2005; Wilson et al., 2017). Test-retest reliability in language production has so far not been studied with MEG, even though the high time resolution of the method makes it particularly well suited for studying the dynamic neural processes underlying language.

We assessed consistent MEG evoked activity in two common tasks, picture naming and semantic judgement. Naming tasks are widely used in studying the process of language production in both clinical populations and healthy participants (Alario et al., 2004; Cornelissen et al., 2003; Laganaro et al., 2015; Liljeström et al., 2009). Naming a depicted item is suggested to incorporate all processing stages that are required for word production, ranging from recognizing the item to be named, to selecting an appropriate concept and word form, and finally to articulation (Indefrey and Levelt, 2004). Previous MEG studies have systematically shown that naming an object induces evoked activity in the occipital cortex within 200 ms of picture presentation, followed by activity in parietal and temporal areas after 200 ms, and activity in frontal and sensorimotor regions after 300–400 ms (Hultén et al., 2009; Indefrey and Levelt, 2004; Liljeström et al., 2009; Salmelin et al., 1994; Sörös et al., 2003; Vihla et al., 2006).

For patients with severe speech difficulties a naming task may be too demanding to perform in a clinical setting. A semantic categorization task (‘Is this item living’) allows for simplified responses (’yes/no’), which might not require as much effort as naming the object, since the response is always the same. Previous MEG studies have shown that a semantic judgement task induces a largely similar evoked activation pattern as picture naming, including involvement of frontal regions (Hultén et al., 2009; Vihla et al., 2006). These results suggest that a semantic judgement task, when assessed using a spoken response, might suffice as a surrogate task to probe the integrity of brain regions related to language production in clinical populations, yet could be easier for the participants to perform (Lupuan and Mirman, 2013).

In addition to the widely used evoked activity, we assessed the consistency of modulations of MEG oscillatory activity related to both picture naming and semantic judgement. Preparation for language production has been shown to modulate cortical oscillatory activity in frontal, motor, temporal and parietal regions (Conner et al., 2014; Flinker et al., 2015; Kojima et al., 2013; Piai et al., 2015; Saarinen et al. 2006) that are typically associated with language processing, but the test-retest consistency of these modulations remains unknown.

## 2 Materials and Methods

### 2.1 Participants

20 healthy human participants participated in the MEG measurements (10 females, 10 males; mean age 25 years; age range 21–35 years). All participants were native Finnish speakers, right-handed, had no history of neurological, psychiatric or developmental disorders or learning disabilities and had normal or corrected-to-normal vision. We obtained a written informed consent from all participants, in agreement with the prior approval of the Aalto University Research Ethics Committee. One participant’s data was not used for the analysis due to non-compliance with the task instruction.

### 2.2 Experimental design

#### 2.2.1 Tasks

The experiment included four tasks: a naming task, a semantic judgement task and two variants of a visual task. In the naming task, the participants were presented with pictures and asked to overtly name the object in the picture. In the semantic task, the participants were asked to overtly say “yes” (“kyllä” in Finnish) if the object in the picture represented a living object and “no” (“ei” in Finnish) if it did not. If the participant did not know the name or the category, he/she was instructed to say “skip” (“ohi” in Finnish). In the visual tasks, participants were asked to say “yes” (“kyllä” in Finnish) when a target picture was presented (picture featuring a red cross in the middle). In one variant of the visual task, the pictures were real identifiable objects and, in the other variant, scrambled objects. In all tasks, the overt response was given after a short delay period. The participants familiarized themselves with the tasks in a practice session before the MEG recording.

#### 2.2.2 Stimuli

As stimuli we used 300 line drawings of objects, and 100 scrambled images constructed from these object pictures. The scrambled images were generated by dividing the original images into 60-by-60 equally sized elements and by randomly permuting 80 % of the locations of the elements that included lines (elements with only white background were kept in place). Scrambled images were also randomly inverted horizontally or vertically.

Half of the stimuli were used in the first measurement session and the other half in the second session. The naming task, semantic task and the visual task with object pictures each used 50 different pictures of objects, which were presented twice, and the visual task with scrambled pictures used 50 different pictures of scrambled objects, which were presented twice. Each visual task, in each measurement session, also included 20 target stimuli, with a red cross in the middle; these stimuli were not included in the analysis.

The naming agreement of the stimulus objects was evaluated by 22 participants (13 females, 9 males; mean age 26 years, age range 19–33 years) who did not participate in the MEG study. At least 16 out of 22 participants used the same name for the 300 objects that were included in the study. The names of the objects had a word length of 3–11 letters (no compounds). The Finnish word frequency value for every object name was derived from a Complete Finnish Wikipedia download (https://dumps.wikipedia.org/fiwiki/); the December 2008 version used here is no longer available online. The cumulative stem frequency value (including all the inflectional variants of a word stem) for every object name was over 0.24 per million words. The categorization consistency (living or nonliving) of the objects was evaluated by another group of nine participants (4 females, 5 males; mean age 26.7 years, age range 20–32 years) who did not participate in the MEG study. In the semantic judgement task, we only used objects for which at least seven out of nine participants agreed on the categorization. There was no significant task difference (Kruskall-Wallis test, p>0.05) in the naming agreement, categorization agreement, word length or word frequency of the object pictures.

As regards the scrambled objects, another set of four participants (2 females, 2 males; mean age 26 years, age range 23–30 years) first named the original objects and then evaluated whether the scrambled images of those objects were identifiable as a certain object. 74 % of the scrambled objects could not be identified by any of these participants. Between all four tasks there was no significant difference in picture luminance (one-way ANOVA, p>0.05).

#### 2.2.3 Stimulus procedure

The picture presentation protocol was the same in all tasks. Pictures (covering 3– 6^°^ × 3–6^°^ of the central visual field) were presented to the participants in blocks consisting of ten pictures (stimulus onset asynchrony, SOA 4.3 s). The visual tasks included 1–3 target pictures per block. Only one task was performed during one block. Before the block began, the task instruction appeared on the screen for 5 s. In the block, a fixation cross first appeared on the screen for 1 s, followed by the picture for 300 ms. After the picture presentation, a blank screen was presented for 1 s, followed by a question mark for 2 s. This sequence was repeated ten times to present the ten pictures in one block. During one block a picture was only presented once (during the overall measurement session, each picture was presented twice). Participants were instructed to give their answer when the question mark appeared on the screen and fixate the centre of the screen during the whole block. See Fig. 1 for an illustration of the experimental design.

**Fig. 1.**
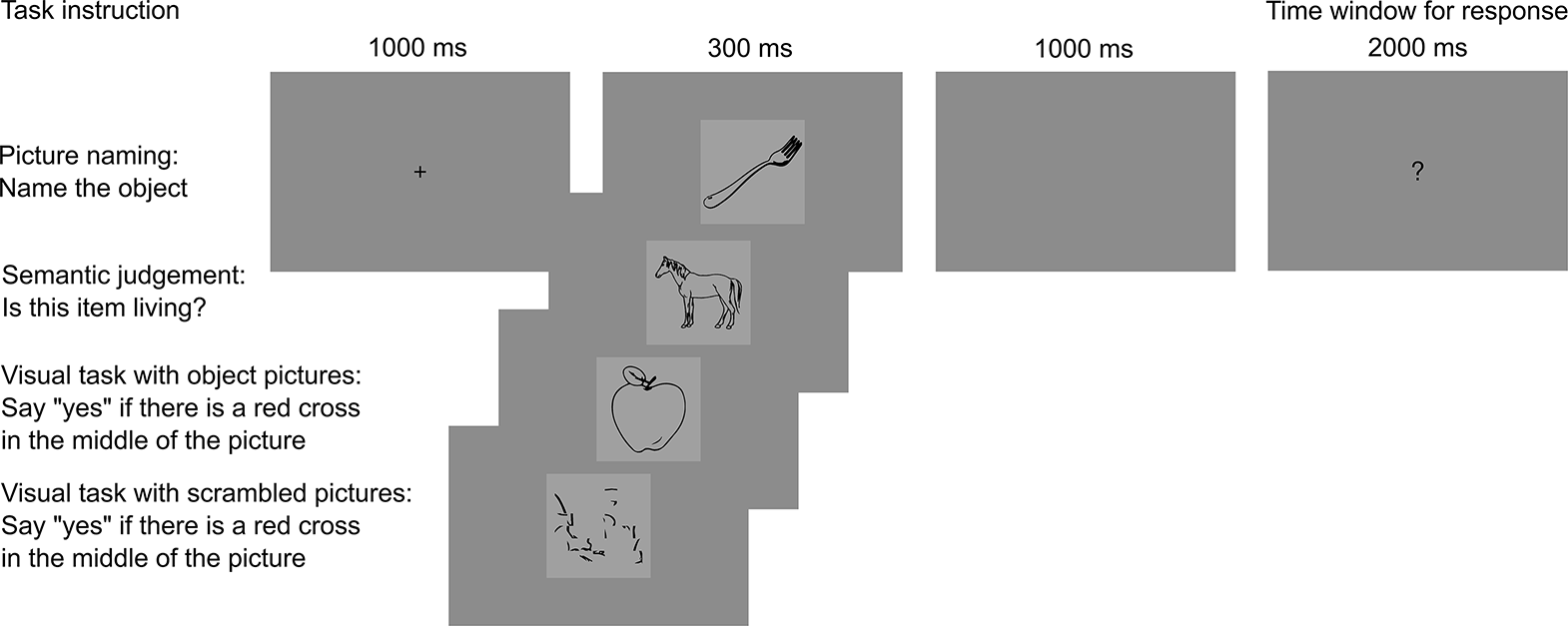
Experimental design. Subjects were asked to perform the task when a question mark appeared.

MEG data was recorded from each participant in two separate measurement sessions, 1–13 days apart. The stimulus pictures were divided into two non-overlapping sets (A, B). Half of the participants started with the set A and the other half with the set B. The study protocol was the same in both measurement sessions. A measurement session included a total of 44 task blocks: 10 blocks each for the naming and semantic tasks and 12 blocks each for the visual tasks. The order of the blocks was randomized. After performing 8–9 blocks, participants had the opportunity for a short break.

In each MEG measurement session we additionally recorded resting data with eyes open and eyes closed. We also recorded MEG data while participants were doing mouth movement and hand tension tasks. Outside the MEG device, participants performed psychological and behavioural tests. These data are not analyzed in the present study.

### 2.3 MEG and MRI recordings

MEG data was measured at the Aalto NeuroImaging MEG Core (Aalto University, Espoo, Finland) in a magnetically shielded room using a Vectorview whole-head MEG device (MEGIN (Elekta Oy), Helsinki, Finland). The device comprises 306 sensors, organized in 102 triplet elements (two planar gradiometers and one magnetometer, each coupled with a Superconducting Quantum Interference Device) in a helmet-shaped array. Vertical and horizontal electro-oculogram (EOG) signals were recorded for identifying eye blinks and saccades. An electromyogram (EMG) was recorded with two electrodes placed side by side near the lower lip margin to monitor mouth movements. The overt speech responses were recorded for later evaluation of correct responses. Five head position indicator (HPI) coils were attached to the head. The positions of these coils with respect to three anatomical landmarks (nasion as well as left and right preauricular points) were digitized using a Polhemus Fastrak (Colchester, VT, US) system. The head position was measured at the beginning of each recording. The MEG signals were filtered at 0.1–330 Hz and sampled at 1000 Hz. MRIs were acquired at the Aalto NeuroImaging Advanced Magnetic Imaging (AMI) Centre with a Siemens Magnetom Skyra 3.0 T MRI scanner using a standard T1-weighted gradient echo sequence.

### 2.4 MEG preprocessing

The spatiotemporal signal space separation method (tSSS; Taulu and Simola, 2006) was applied for noise reduction, with a 16 s temporal window, a subspace correlation limit of 0.98, inside expansion order of 8, and outside expansion order of 3. The measured head positioning data was used to transform the MEG data recordings to a common position (MEGIN (Elekta) Maxfilter software package, 2.2.12). Epochs containing eye blinks or saccades were rejected. The EOG rejection limit was determined separately for each participant (range 100–150 μV). Only epochs corresponding to correct responses were analysed; synonyms were considered as correct responses in the naming task. If the data from a participant yielded less than 70/100 accepted trials after exclusion of epochs with incorrect responses or blinks, even in one task condition, independent-component-analysis (ICA) based projection of eye-blink artefacts was applied to that participant’s whole MEG data set in that measurement session to enhance the number of artefact-free trials. In this case, EOG-based rejections were not applied.

### 2.5 Analysis of evoked activity

MEG data was low-pass filtered at 40 Hz and averaged over epochs from 200 ms before to 1000 ms after the stimulus onset, separately for each task. Epochs were baseline-corrected to the 200-ms interval preceding the stimulus onset. A source-level overview of the spatiotemporal distribution of neural activity was obtained with cortically constrained L2 minimum-norm estimates (MNEs) using MNE Python version 0.8 (Gramfort et al., 2013). The cortical surface was reconstructed for each participant from the MR images using the Freesurfer software package (Dale et al., 1999, Fischl et al., 1999). A surface-based cortical source space (average spacing between equivalent current dipoles 9.9 mm, not including the cerebellum) was used. A loose orientation constraint of 0.3 was applied such that currents normal to the cortical surface were favoured by reducing the variance of the transverse source components. Depth weighting of 0.8 was used to reduce the inherent bias of the MNE towards superficial sources. A single-layer boundary-element model (BEM) was determined from the inner skull surface and used as a head conductor model in the forward computation. A noise covariance matrix was estimated from the unaveraged 200-ms pre-stimulus baseline periods of all four tasks for each measurement session. The noise covariance matrix regularization factor for magnetometers and gradiometers was 0.05. Noise-normalized MNEs (dynamical Statistical Parametric Maps, dSPMs) were calculated over the whole cortical area to estimate the signal-to-noise ratio at each potential source location. For further analyses, the individual MNEs were morphed, with spatial smoothing, to a standard template brain (fsaverage as provided by FreeSurfer).

Evoked response analyses were performed for selected time windows (0–200 ms, 200– 400 ms, 400–600 ms and 600–800 ms) time-locked to stimulus presentation, based on a neurocognitive model of speech production (Indefrey and Levelt, 2004) and earlier MEG findings (Hultén et al., 2009; Liljeström et al., 2009; Salmelin et al., 1994; Sörös et al., 2003; Vihla et al., 2006).

### 2.6 Analysis of cortical oscillatory activity

Task-related modulations in the power level of cortical oscillations were estimated using a linearly constrained minimum variance spatial filter (beamformer), event-related Dynamic Imaging of Coherent Sources (erDICS; Laaksonen et al., 2008), applying in-house Matlab scripts. The time-frequency representation for each epoch, from −400 ms to 1200 ms with respect to the stimulus onset, was first calculated using Morlet wavelets of width 7, at the sensor level. The cross-spectral density (CSD) matrix for every task in each measurement session was obtained by calculating the product of the time-frequency representations of the epoch time series for all sensor combinations. The single-trial CSDs were then averaged into a mean CSD matrix for every task in each measurement session. The analysis was performed for frequencies ranging from 4 to 90 Hz and averaged across pre-specified frequency bands (theta 4–7 Hz, alpha 8– 13 Hz, low beta 14–20 Hz, high beta 21–30 Hz, low gamma 31–45 Hz, high gamma 60–90 Hz), adapted from Liljeström et al., 2015a.

The sensor-level data was transformed into a cortical representation using the DICS spatial filter (Gross et al., 2001). The source space was the same as for the cortically constrained MNEs (see analysis of evoked activity above). The source space (maximum distance to the sensors 7 cm, approximately 700 vertices per hemisphere) was defined for every participant based on a reference participant (one of the study participants). A common spatial filter across tasks was constructed for each cortical source point in the frequency bands and time windows of interest. Estimates for the power level of cortical oscillations for every task in each measurement session were obtained by applying this spatial filter and selecting the source orientation that yielded the maximum power.

Source-level power estimates were normalized for further analyses. The standard deviation of the power estimates across all task conditions and all source points was calculated and the power estimates in each task condition and at each vertex were divided by this standard deviation. This normalization was done separately for every participant, time window, frequency band and measurement session.

For analysis of cortical oscillatory activity, we divided the time interval from the onset of the image to the (delayed) request to give the overt response into three separate time windows: 0–400 ms, 400–800 ms, and 800–1200 ms. We based this choice on a previous study (Laaksonen et al., 2012) that showed modulations of cortical oscillations preceding onset of speech and illustrated how oscillatory modulation typically lasts longer than evoked activity.

### 2.7 Consistency analysis

#### 2.7.1 Parcellation

A custom-made parcellation based on the Destrieux FreeSurfer template parcellation for fsaverage was used. The custom-made parcellation was constructed using a merge- and-split approach to produce uniform-sized parcels, while preserving coarse anatomical boundaries when possible. First, gyri and sulci belonging to the same structure (e.g. middle temporal gyrus and sulcus) were merged. This resulted in elongated regions that, using principal component analysis (PCA), were split in the direction perpendicular to the largest principal component into parcels with similar size across the entire cortex. In this study, a parcellation including 55 brain regions per hemisphere was used.

#### 2.7.2 Task-related modulation of evoked activity and oscillatory activity

To identify naming-related changes of evoked activity and modulation of cortical oscillatory activity we compared the naming task with the visual task with object images. A similar procedure was used to define semantic judgement-related activity. We selected the visual task with object pictures as a control condition to rule out the possible activations related to object recognition (see Price et al., 2005). Statistical analyses (repeated measures t-test) were done at the brain-region level, separately for each time window, frequency band and measurement session. For the subsequent analysis of consistency, we selected the task-related evoked activity and modulation of oscillatory activity that was statistically significant (p<0.005, uncorrected) in both measurement sessions.

#### 2.7.3 ICC analysis

We used intraclass correlation coefficient (ICC) analysis (Shrout and Fleiss, 1979) to quantify the test-retest reliability of the task-related modulation of evoked and oscillatory activity. ICC determines the consistency of activation in individual subjects across days by evaluating whether the rank order of individual activations is preserved across the days. ICC has been the measure of choice in recent neuroimaging studies that examined test-retest reliability (see e.g. Cao et al., 2014; Lee et al., 2010; Martin-Buro et al., 2016; Plichta et al., 2012; Vetter et al., 2017). ICC analyses were done for the predefined (see section: 2.7.2 Task-related modulation of evoked activity and oscillatory activity) task-related activity (difference of source-level estimates between tasks) separately for each brain region, time window and frequency band. ICC can be formulated as:

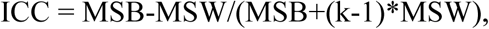

Where MSB = mean square between-subjects, MSW = mean square within-subjects, and k = number of repeated sessions. ICC increases with increasing between-subject variance and with decreasing within-subject variance. We denote ICC between 0.40 and 0.59 as fair, between 0.60 and 0.74 as good and between 0.75 and 1.00 as excellent consistency (Cicchetti, 1994).

### 2.8 Data and code availability

The raw MEG data cannot be made openly available, according to the ethical permission and national privacy regulations at the time of the study. The derived anonymous results that served as input for the figures will be made openly available.

Analysis methods used in the study are based on openly available software packages for MEG analysis, as described in the section 2 Methods. Methods that were used for analysis of cortical oscillations were incorporated into MNE Python version 0.16.

## 3. Results

We determined whether MEG evoked activity in a picture naming paradigm is consistent across measurement days. We also determined whether a semantic judgement task elicits similarly consistent activation patterns as the naming task. Furthermore, we investigated whether modulation of cortical oscillatory activity occurs consistently in brain regions previously shown to be engaged in language production.

### 3.1 Consistent evoked activity in picture naming

The typical activation patterns related to picture naming that have been reported earlier (Hultén et al., 2009; Liljeström et al., 2009; Vihla et al., 2006) were observed also in the present study (Fig. 2). The group-level analysis revealed an activation pattern that encompassed perisylvian language regions from 200 ms onwards, including the middle temporal cortex and frontal cortex from 400 ms onwards, on both measurement days. As illustrated in Fig. 3, the left-hemisphere activation pattern showed fair to excellent consistency across the measurement days. Consistent evoked activity was detected in the left frontal (400–800 ms), left sensorimotor (200–800 ms), left parietal (200–600 ms), left temporal (200–800 ms), left occipital (400–800 ms) and left cingulate (600– 800 ms) regions, as well as in the right superior temporal (600–800 ms) region. No consistent evoked activity was detected in the early 0–200 ms time window. Figure 4 focuses on the consistent activity at 400–600 ms to illustrate the individual-level data. The strength of the evoked activity (difference between picture naming and visual task) is shown for each subject in ten brain region that showed consistent activation for the two different measurement sessions. This evoked activity was positive in most individuals, indicating larger activation for picture naming compared to the visual task. As shown in Fig. 4, the variance within subjects (MSW) was small in comparison to variance between subjects (MSB). Significant Pearson’s correlation between measurement days is in line with the ICC analysis. See Supplementary Fig. S1 for individual activations on day 1 and day 2 at 400–600 ms.

**Fig. 2.**
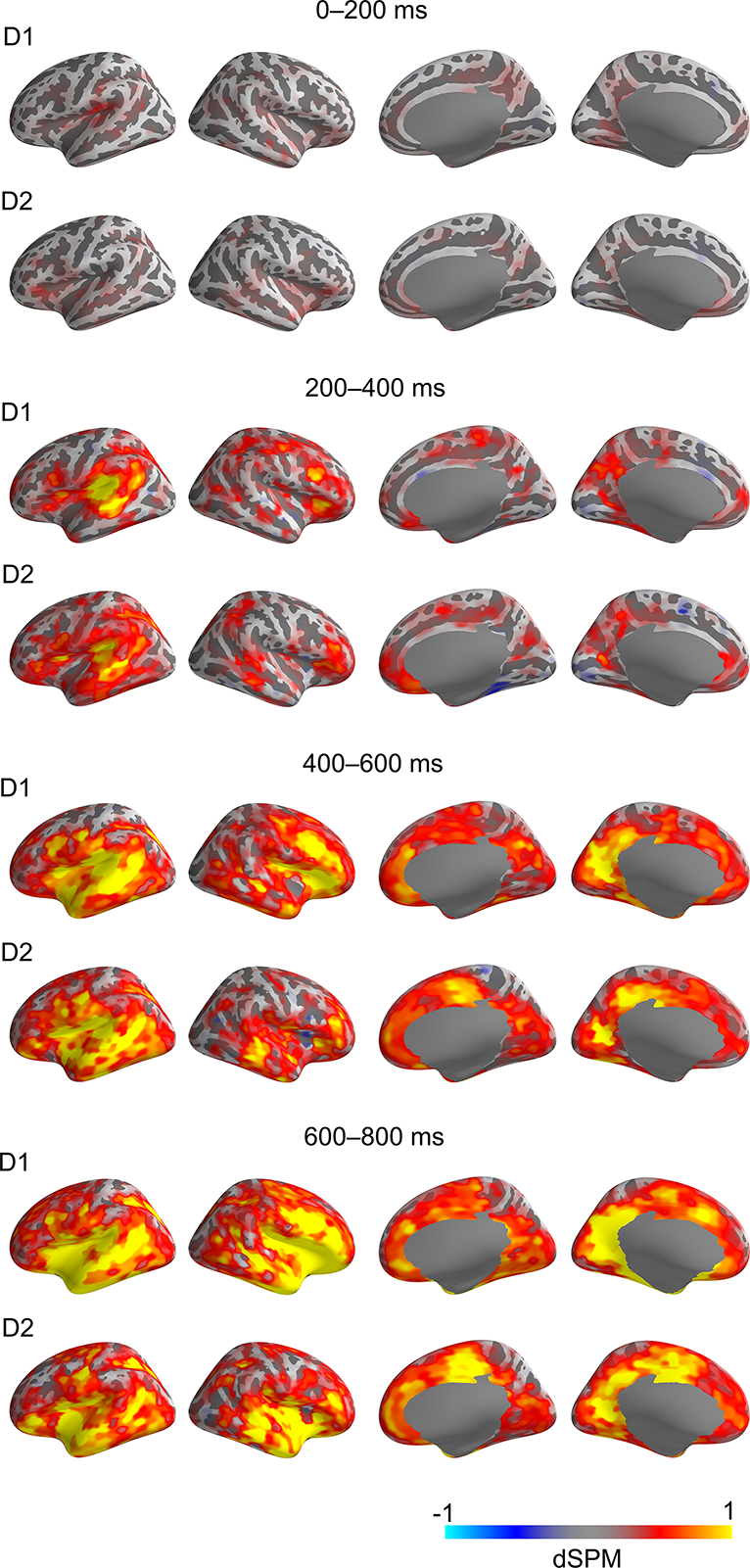
Overall activation pattern in picture naming in different time windows after stimulus onset. For each time window, the evoked response strength in the naming task, as compared to the visual task, on day 1 (D1) and day 2 (D2) is shown. The color scale indicates the difference in dSPM values between the two tasks (yellow/red colors indicate stronger responses to naming; for significance testing, see Fig 3.) Each row depicts activation strength in left lateral, right lateral, right medial and left medial cortex.

**Fig. 3.**
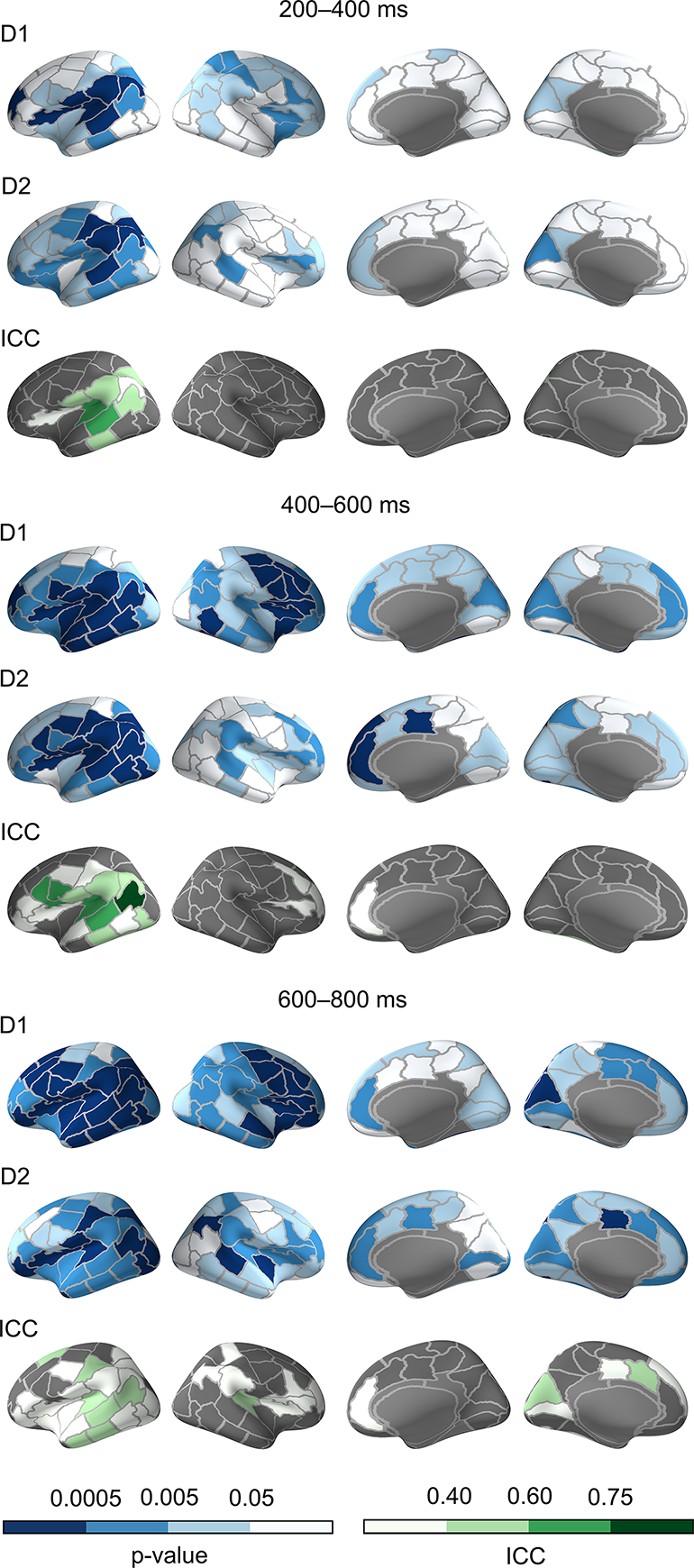
Consistent evoked activity in picture naming in different time windows after stimulus onset. Evoked activity related to picture naming on day 1 (D1) and day 2 (D2) are illustrated at different p-values (dark blue, p<0.0005, mid blue, p<0.005, light blue p<0,05, white n.s.). Grey parcels were not incorporated into the ICC analysis as they did not meet our significance criteria (p-value <0.005 on both days). Consistent evoked activity (ICC) is shown with excellent (ICC>0.75, dark green), good (ICC 0.6–0.75, mid green), fair (ICC 0.4–0.6, light green), or poor consistency (ICC<0.4, white). Left lateral, right lateral, right medial and left medial cortex are shown.

**Fig. 4.**
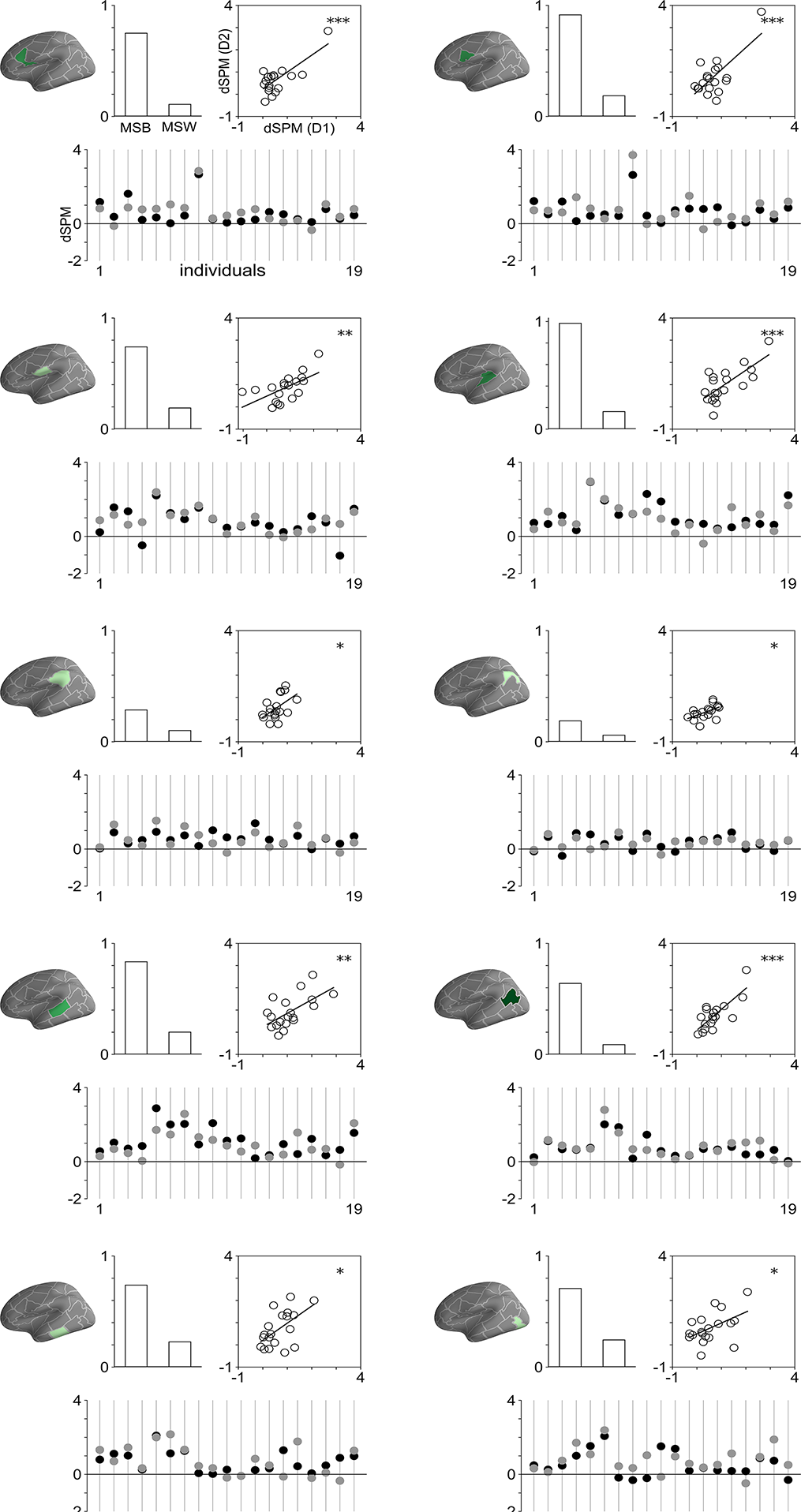
Evoked activity of individual subjects for picture naming in the time window 400–600 ms in ten regions that showed consistent naming-related evoked activity. The lower graph, for each brain region, presents the evoked response strength (dSPM) in the picture naming task compared to the visual task, for all the 19 participants (Day 1 black; Day 2 grey). The scatter plot shows the above-mentioned individual data plotted for day 1 (D1) and day 2 (D2). The significance of the Pearson correlation is indicated (p<0.05^*^, p<0.01^**^, p<0.001^***^). The bar graph shows between-subject variance (MSB) and within-subject variance (MSW). See left upper corner for units.

### 3.2 Consistent evoked activity in semantic judgement

In line with previous studies (Hultén et al., 2009; Vihla et al., 2006), the activation pattern elicited by semantic judgement encompassed brain regions within the perisylvian language regions, bilaterally, as indicated by difference in activation between the semantic judgment and the visual task (Fig. S2). However, fairly few brain regions showed a significant difference of at least p<0.005 (uncorrected) between conditions on both days (Fig. 5; all non-grey regions). Amongst those regions, good or excellent consistency (Fig. 5; mid to dark green color) was found in the left mid-part of the superior temporal cortex (200–800 ms,). Consistency (Fig. 5) was also found within the left sensorimotor cortex (400–800 ms), the mid-part of the left inferior temporal cortex (600–800 ms), left subparietal cortex (600–800 ms), left occipital cortex (400– 600 ms) and right supramarginal cortex (600–800 ms). No consistent evoked activity was detected in the early 0–200 ms time-window.

**Fig. 5.**
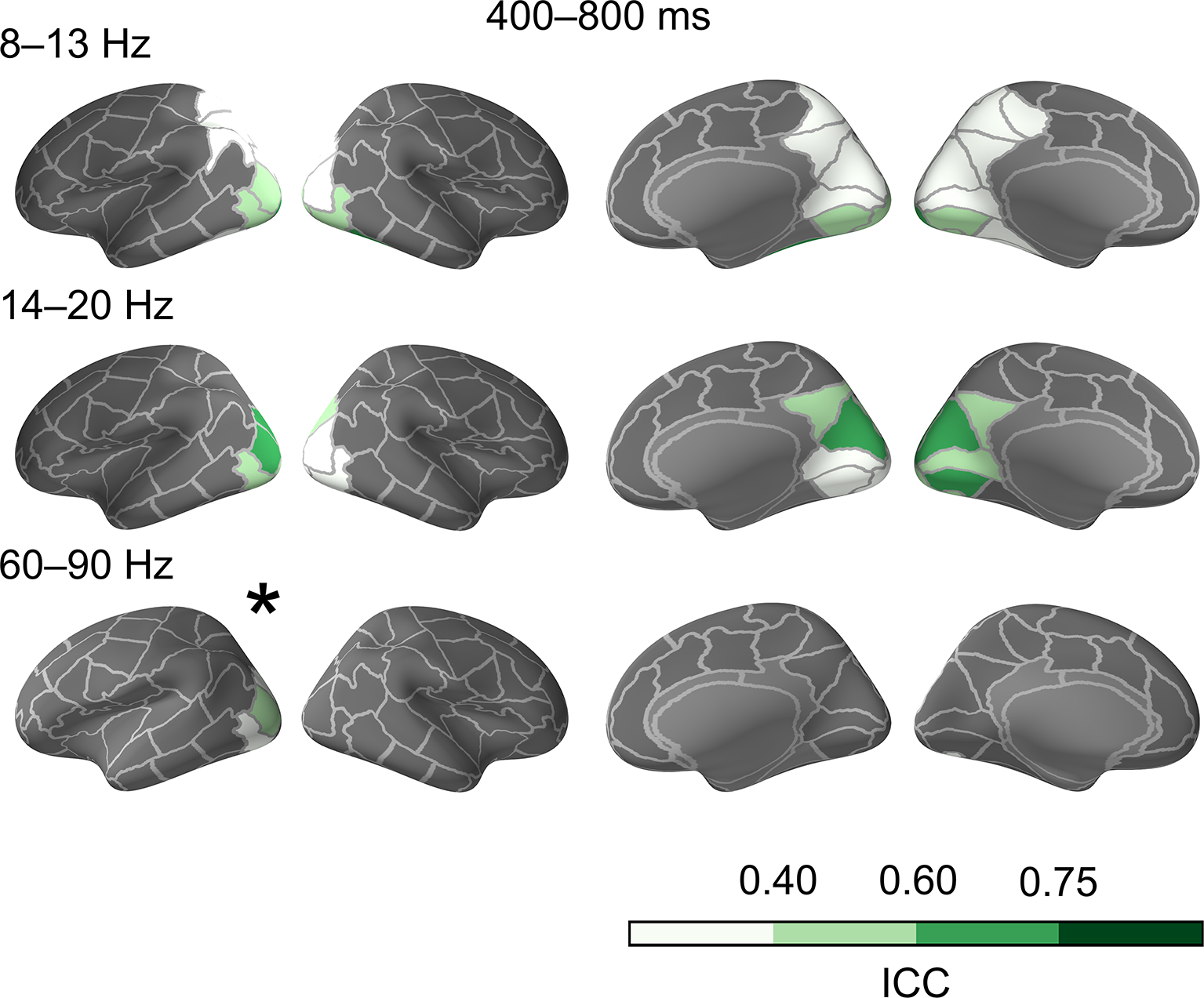
Consistent evoked activity (ICC) for semantic judgements in different time windows after stimulus onset. Non-grey parcels represent the brain regions that were selected for the ICC analysis, with the different shades of green denoting excellent (ICC > 0.75, dark green), good (ICC 0.6–0.75, mid green) or fair (ICC 0.4–0.6, light green) consistency. Left lateral, right lateral, right medial and left medial cortex are shown.

### 3.3 Consistent modulation of cortical oscillatory activity in picture naming

Consistent suppression of cortical power in picture naming was predominantly observed bilaterally in posterior brain areas, including occipital, cingulate, occipitotemporal, and parietal cortex in the 4–7 Hz, 8–13 Hz and 14–20 Hz frequency bands from 400 ms onwards (Fig. 6). However, consistent suppression of power was also observed in motor regions in the 14–20 Hz and 21–30 Hz bands, particularly within the time window 800–1200 ms (see also Supplementary Fig. S3). Furthermore, power enhancement was observed at 14–20 Hz and 21–30 Hz in the right temporal region (0– 400 ms). Consistent power enhancement within the 60–90 Hz band was restricted to occipitotemporal cortex.

**Fig. 6.**
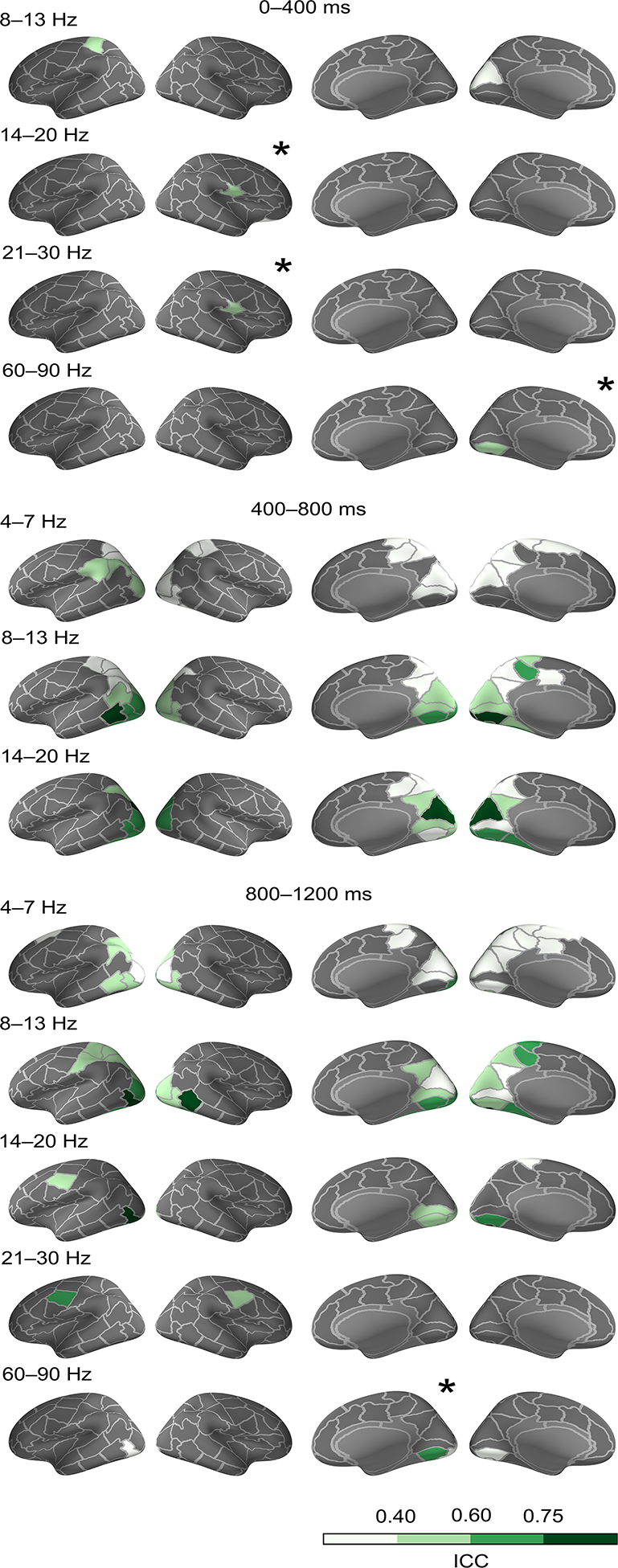
Consistent modulation of cortical oscillatory activity in picture naming in different time windows after stimulus onset. The observed modulation was a suppression of power, except for activity marked with (*), which represents enhancement of power. Non-grey parcels represent the brain regions that were selected for the ICC analysis. Consistency is denoted as excellent (ICC > 0.75, dark green), good (ICC 0.6–0.75, mid green) or fair (ICC 0.4–0.6, light green). Left lateral, right lateral, right medial and left medial cortex are shown.

### 3.4 Consistent modulation of cortical oscillatory activity in semantic judgement

In semantic judgement, consistent modulation of cortical spectral power was limited to bilateral posterior areas in the occipital, occipitotemporal, and subparietal regions, and the time window 400–800 ms (Fig. 7). In this time window, power was suppressed in 8–13 Hz and 14-20 Hz bands and power was enhanced at 60–90 Hz. See supplementary figure (Fig. S4) of low beta (14–20 Hz) power modulation on day 1 and day 2.

**Fig. 7.**
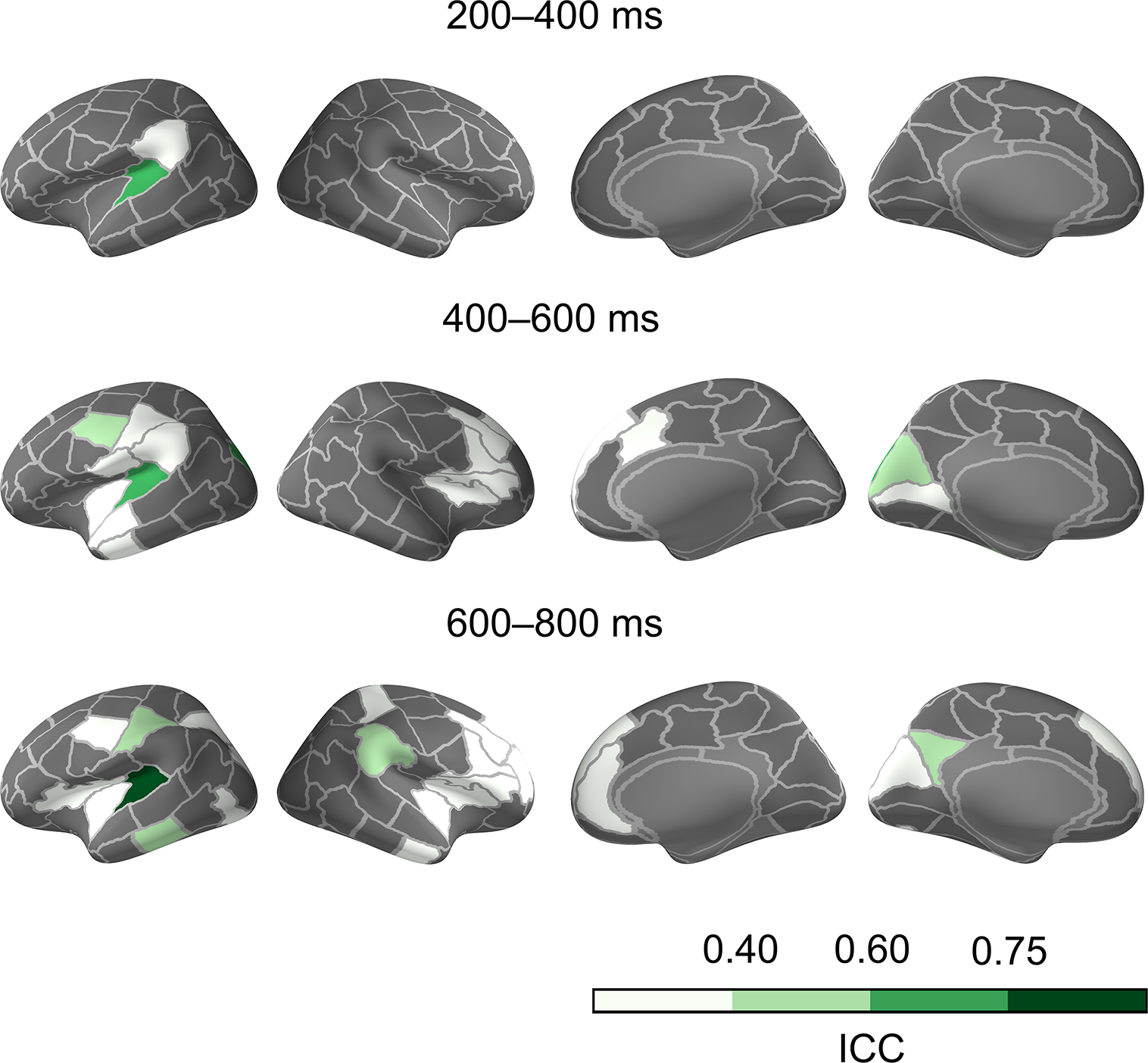
Consistent modulation of cortical oscillatory activity in semantic judgement in different time windows after stimulus onset. Modulation was observed as a suppression of power, except in brain regions marked with (^*^), where enhancement was observed. Non-grey parcels represent the brain regions that were selected for the ICC analysis. Consistency is denoted as excellent (ICC > 0.75, dark green), good (ICC 0.6–0.75, mid green) or fair (ICC 0.4–0.6, light green). Left lateral, right lateral, right medial and left medial cortex are shown.

## 4 Discussion

Reliable measures of brain activity related to language production in individual subjects are essential for both basic research and, in particular, for clinical applications of brain imaging methods. Here, consistency was assessed for cortical activation patterns related to picture naming and semantic judgement. From the MEG recordings, we determined modulations of both evoked and oscillatory activity that were consistent across measurement days in healthy individuals. In picture naming, we found consistent left-lateralized evoked activity patterns in the parietal and temporal regions starting at 200 ms after picture presentation, followed by activity in the frontal cortex from 400 ms onwards. In semantic judgement, highly consistent activation was restricted to the left superior temporal cortex. Consistent modulation of oscillatory activity was mainly identified in posterior cortical regions. In addition, for the naming task we detected typical beta oscillatory suppression in the motor region from 800 ms onwards.

### 4.1 Picture naming evoked responses yield consistent left-lateralized activation patterns

This MEG study demonstrated the consistency of the left-hemisphere spatiotemporal sequence of activation described in previous MEG studies and neurocognitive models of picture naming (Hultén et al., 2009; Indefrey and Levelt, 2004; Liljeström et al., 2009; Salmelin et al., 1994; Sörös et al., 2003; Vihla et al., 2006): we found consistent evoked activity in the left parietal (200–600 ms), temporal (200–800 ms), frontal (400– 800 ms), sensorimotor (200–800 ms), occipital (400–800 ms) and cingulate (600–800 ms) regions, and in the right superior temporal (600–800 ms) cortex. Activation in parietal and temporal areas after 200 ms has been suggested to represent identification and semantic processing of an object (Clarke et al., 2015; Sudre et al., 2012). Activation in the frontal and sensorimotor regions after 300 ms has been suggested to be related to phonological processing and motor preparation for articulation (Indefrey and Levelt, 2004; Vihla et al., 2006). The consistent evoked spatiotemporal activity pattern related to picture naming thus seemed to capture the full set of cortical areas related to preparation for language production, including the inferior frontal gyrus. This would be a great advantage since previous fMRI studies on the reproducibility in language production have suggested that in order to reliably capture the whole language production related brain network, multiple different language tasks should be used and combined (Harrington et al., 2006; Nettekoven et al., 2018; Rau et al., 2007).

Previous fMRI reliability studies have shown variable results on the consistency of different brain areas in language production (see Eaton et al., 2008; Gorgolewski et al., 2013; Harrington et al., 2006; Meltzer et al., 2009; Morrison et al., 2016; Nettekoven et al., 2018; Rau et al., 2007; Rutten et al., 2002; Wilson et al., 2017). For example, there have been contradictory findings on whether the IFG, often thought to be crucial for speech production, is consistently activated in speech production tasks (Eaton et al., 2008; Harrington et al., 2006; Meltzer et al., 2009; Nettekoven et al., 2018; Rau et al., 2007). The IFG has been extensively studied in clinical aphasia: better performance in speech production tasks and better recovery are related to greater left IFG activation (Saur et al., 2006; Turkeltaub et al., 2011; Winhuisen et al., 2005). For that reason, brain stimulation rehabilitation protocols aiming to improve language production often seek to activate the left IFG (Holland et al., 2011) and inhibit the right IFG (Naeser et al., 2005; Saur et al., 2006; Turkeltaub et al., 2011).

The present MEG study shows that consistent evoked activity can be detected in the left IFG during a picture naming task. Furthermore, the results provide spatiotemporal information of the left IFG activity. Firstly, consistent naming-related evoked activity was found in the posterior part of the IFG. This observation is in line with several studies emphasizing the role of the posterior part of the left IFG in language production and, in particular, in its core function in phonological processing (reviewed in Costafreda et al., 2006). Secondly, our results show that the consistent naming-related evoked activity in the left posterior IFG occurs at 400–600 ms after stimulus presentation. Phonological processing in the IFG has been suggested to take place in this time window (Indefrey and Levelt, 2004; Sahin et al., 2009). Therefore, we propose that the consistent left IFG naming-related evoked activity from 400 to 600 ms after stimulus presentation observed here is tightly linked to preparation for language production. We further suggest that this time window should be used in clinical protocols when targeting left IFG activity related to language production.

### 4.2 Semantic judgement evokes consistent activity in the left temporal cortex

One aim of this study was to determine whether a semantic judgement task with a spoken response (‘yes/no’) would suffice to induce reliable activity in brain regions related to language production and thus offer an option for use on patients with naming difficulties. Our findings suggest that this is not the case. Consistent evoked activity in the semantic judgement was restricted to the left temporal, sensorimotor, subparietal, occipital and right supramarginal cortex. The reason for this spatially limited consistency can be that the difference in activation strength in the semantic judgement task compared to visual task was small; task-related activity was seen in fairly few brain regions. It has been shown that activations with magnitude close to the noise level are less reliable (Meltzer et al., 2009). The reason for the lack of activation can be that the semantic task might not require enough phonological effort since the response is always the same (’yes/no’). Functional imaging studies suggest that low-effort speech tasks do not engage the language system to the extent that more complex language tasks do (Bookheimer et al., 2000; Meltzer et al., 2009; Vanlancker-Sidtis et al., 2003). However, amongst the fairly few consistently activated brain regions, highly consistent activity in the left superior temporal cortex was observed, indicating its relevance for performing semantic judgements. We also detected left sensorimotor activity from 400 ms onwards, suggesting that the semantic judgement task captured activity related to motor preparation for articulation of the formulaic ‘yes/no’ responses.

### 4.3 Cortical oscillatory activity in motor regions is consistently modulated in picture naming

As regards modulation of oscillatory activity in naming, we demonstrate consistent alpha and beta suppression, albeit limited to fairly few brain regions: sensorimotor, parietal and posterior temporal cortex. This is in line with previous studies of power spectral modulation in language production (Laaksonen et al., 2012; Piai et al., 2015). For example, we demonstrate beta (14–20 Hz and 21–30 Hz) oscillatory suppression in the motor regions from 800 ms onwards. We also illustrate consistent occipital and occipitotemporal suppression of alpha (8–13 Hz) and beta (14–20 Hz) bands in both picture naming and semantic judgement tasks, likely related to attentional processing of visual stimuli (Sauseng et al., 2005; Waldhauser et al., 2012).

### 4.4 Conclusions

Overall, our findings highlight evoked activity in picture naming as the most consistent functional marker of the extended cortical system that supports language production in individual participants. The present study emphasizes the usability of the naming task in clinical protocols and demonstrates the relevance of evaluating both group-level and individual-level consistency in neuroimaging studies of human cognition.

## Acknowledgments

This work was supported by Jenny and Antti Wihuri Foundation; Swedish Cultural Foundation in Finland; Maud Kuistila Foundation; Academy of Finland [grant number 315553]; and The Sigrid Juselius Foundation. We thank D. Sc. Tiina Lindh-Knuutila for computing the Finnish word frequencies for object names used in this study.

---

Fig. S1. For every individual the evoked response strength in the picture naming task, as compared to the visual task, on day 1 (upper row) and day 2 (lower row) at 400–600 ms. The color scale indicates the difference (dSPM) between the two tasks (yellow/red colors indicate stronger responses to naming). Left and right lateral cortices are shown.

Fig. S2. Overall activation pattern for semantic judgement in different time windows after stimulus onset. For each time window, the evoked response strength in the semantic judgement task, as compared to visual task, on day 1 (D1) and day 2 (D2) is shown. The color scale indicates the difference (dSPM) between the two tasks (yellow/red colors indicate stronger responses to semantic judgement). Each row depicts activation strength in left lateral, right lateral, right medial and left medial cortex.

Fig. S3. Low beta (14-20 Hz) power modulation in the picture naming task in different time windows after stimulus onset. For each time window, power modulation in the picture naming task, as compared to the visual task, on day 1 (D1) and day 2 (D2) is shown. Yellow/red colors indicate power enhancement and blue colors indicate power suppression. Left lateral, right lateral, right medial and left medial cortex are shown.

Fig. S4. Low beta (14-20 Hz) power modulation in the semantic judgement task in different time windows after stimulus onset. For each time window, power modulation is depicted in the semantic judgement task, as compared to the visual task, on day 1 (D1) and day 2 (D2). Yellow/red colors indicate power enhancement and blue colors indicate power suppression. Left lateral, right lateral, right medial and left medial cortex are shown.

