## Supplementary figures and images for "Picture naming yields highly consistent cortical activation patterns: test-retest reliability of magnetoencephalography recordings"

### Fig. S1.

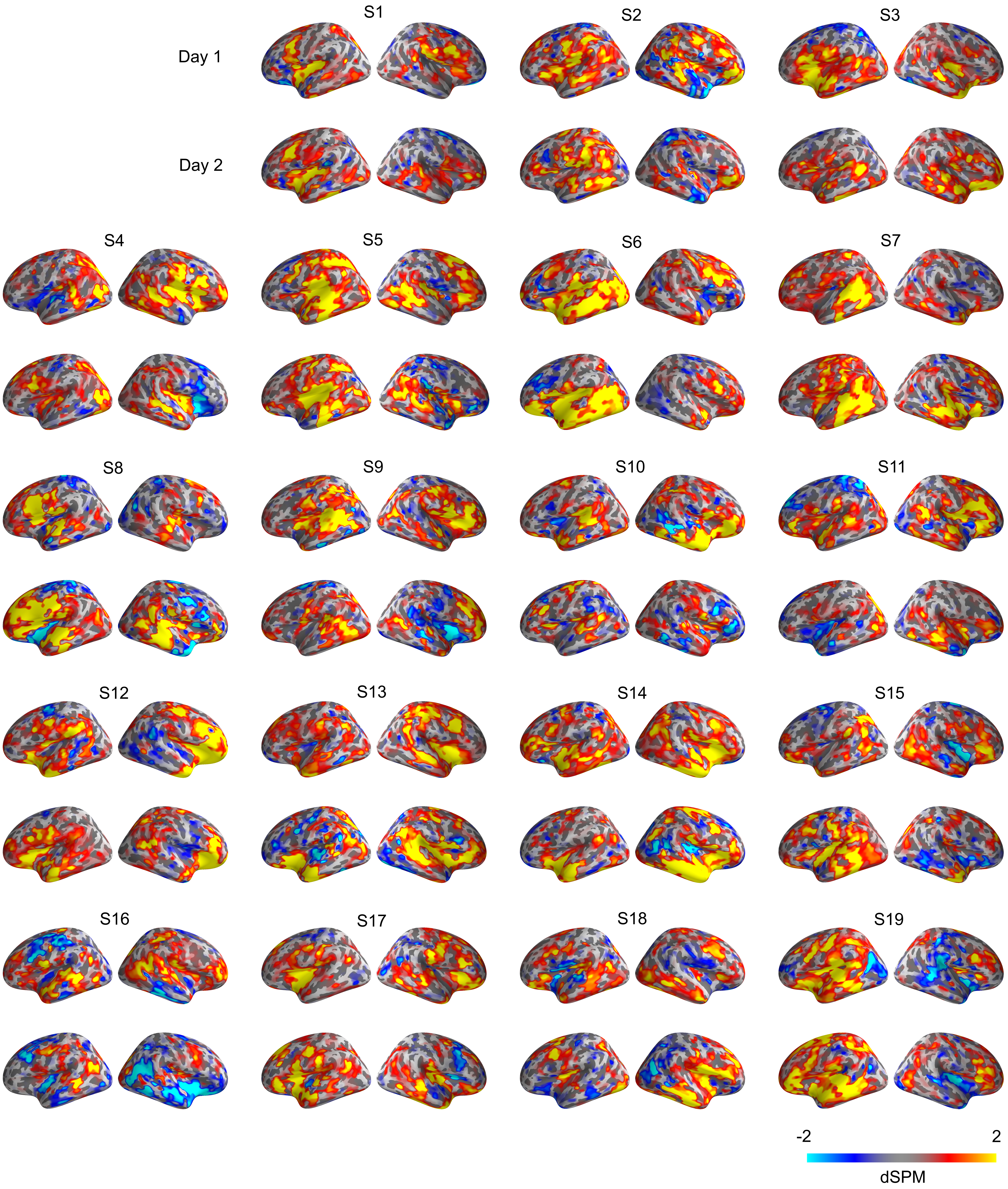

### Fig. S2.

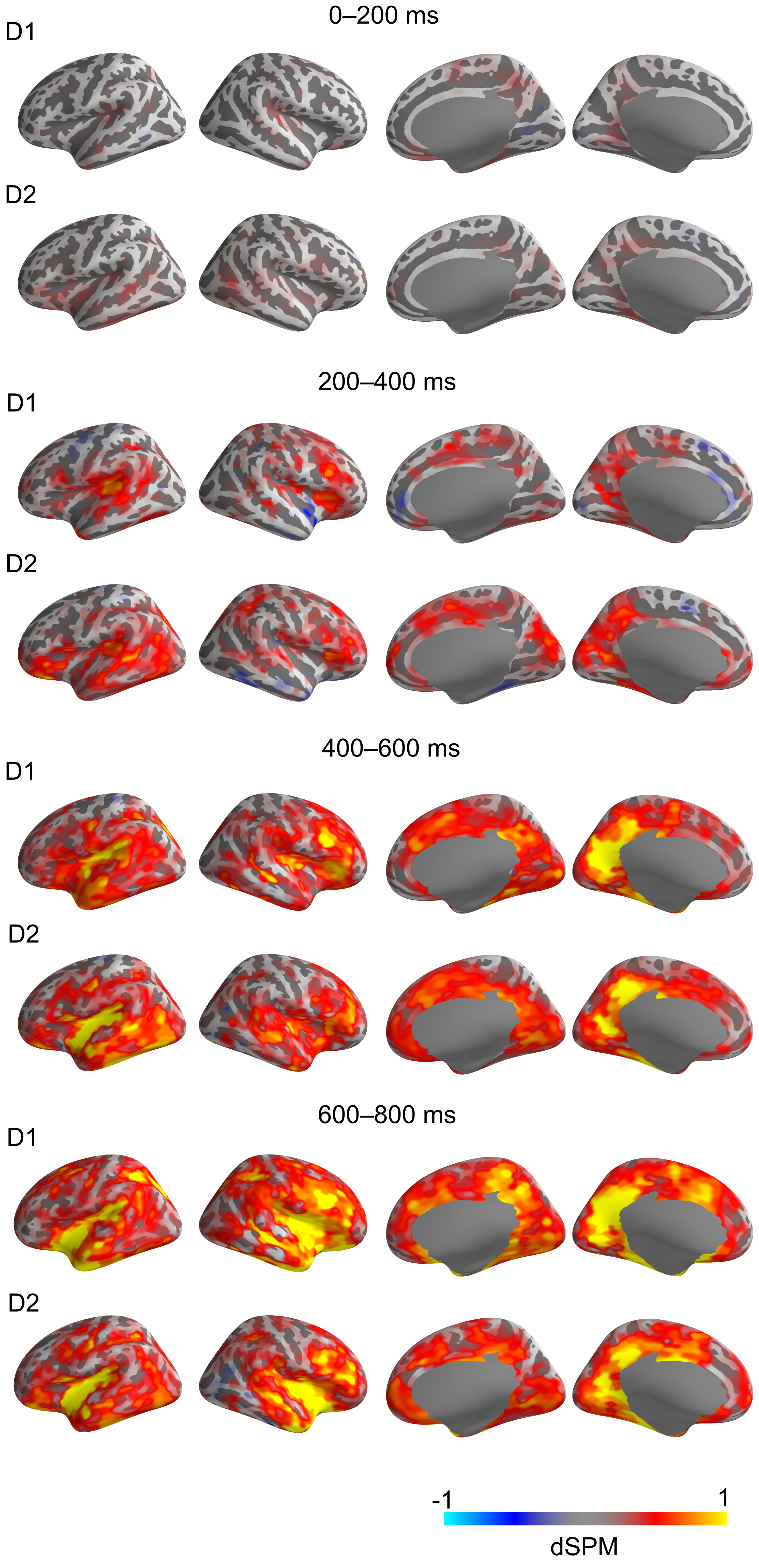

### Fig. S3.

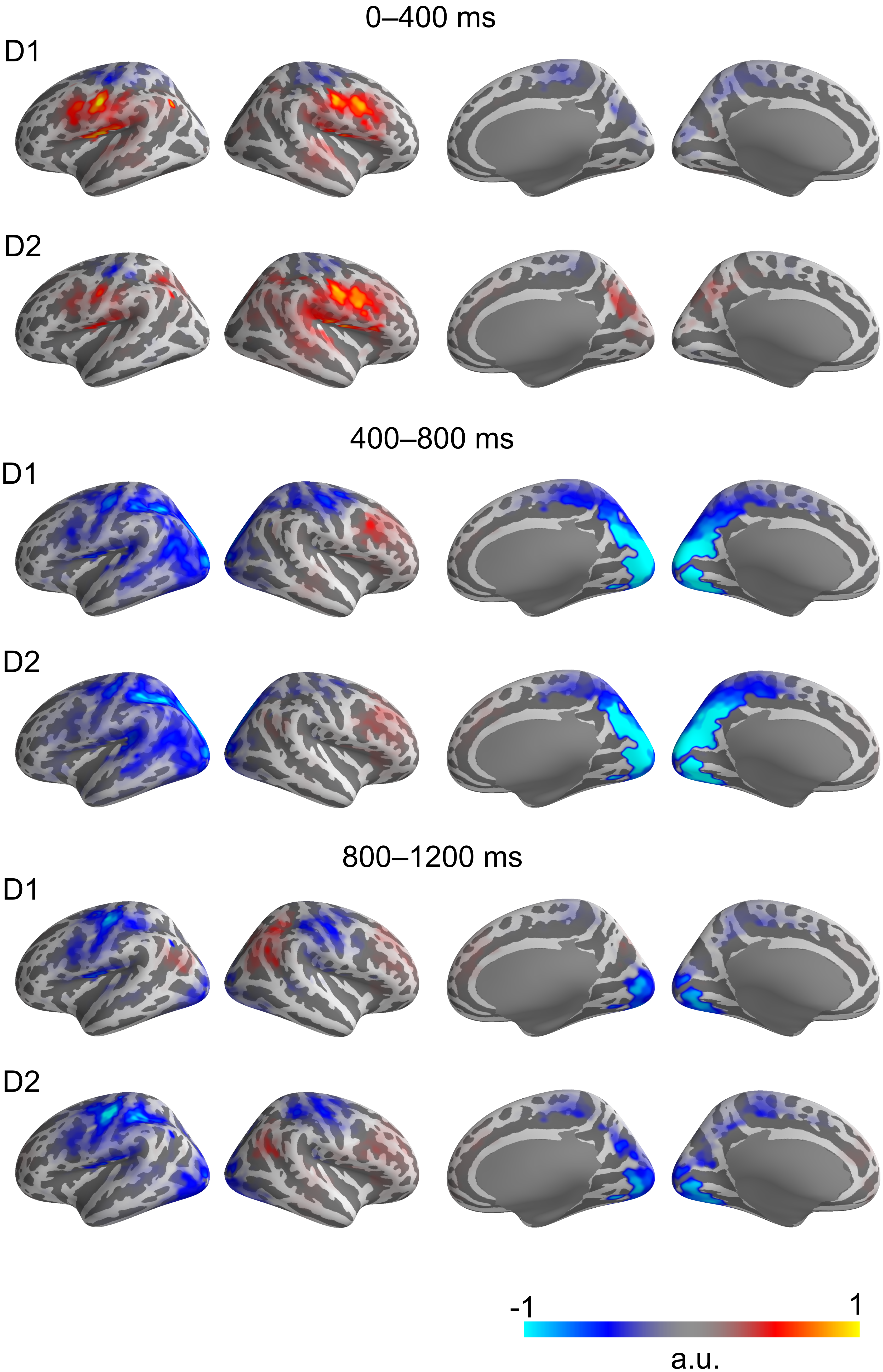

### Fig. S4.

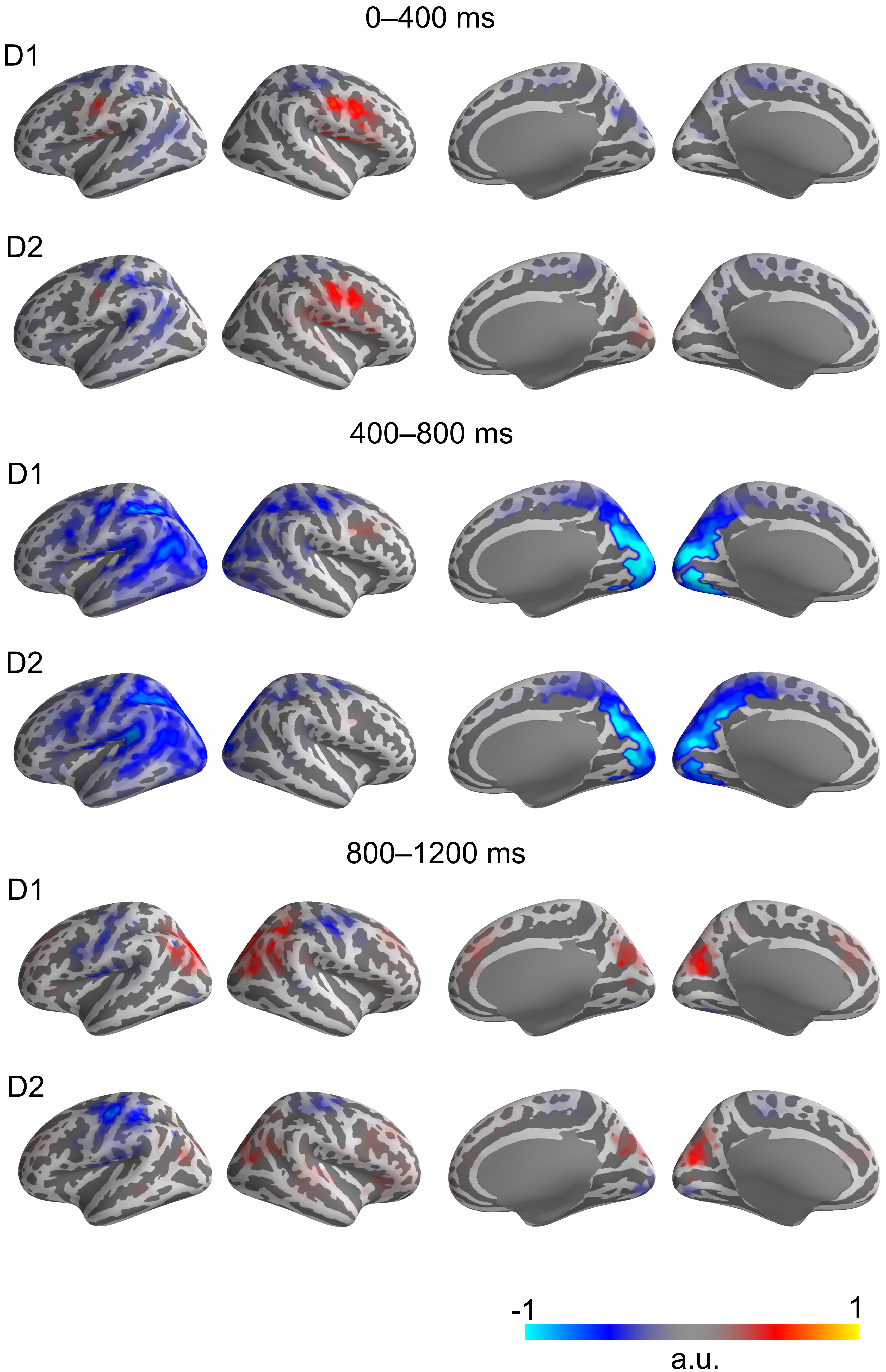
